# A glimpse of antimicrobial resistance gene diversity in kefir and yoghurt

**DOI:** 10.1101/2020.09.02.279877

**Authors:** Adrienn Gréta Tóth, István Csabai, Gergely Maróti, Ákos Jerzsele, Attila Dubecz, Árpád V. Patai, Sára Ágnes Nagy, László Makrai, Krisztián Bányai, Géza Szita, Norbert Solymosi

**Affiliations:** University of Veterinary Medicine Budapest, Centre for Bioinformatics, Budapest, 1078, Hungary; University of Veterinary Medicine Budapest, Department of Microbiology and Infectious Diseases, Budapest, 1143, Hungary; Eötvös Loránd University, Department of Phyisics of Complex Systems, Budapest, 1117, Hungary; Institute of Plant Biology, Biological Research Center, 6726 Szeged, Hungary; University of Veterinary Medicine Budapest, Department of Pharmacology and Toxicology, Budapest, 1078, Hungary; Department of Surgery, Paracelsus Medical University, Nuremberg, Germany; Semmelweis University, 1st Department of Surgery and Interventional Gastroenterology, Budapest, 1082, Hungary; Semmelweis University, Interdisciplinary Gastroenterology (IGA) Working Group, Budapest, 1085, Hungary; Institute for Veterinary Medical Research, Centre for Agricultural Research, Budapest, 1143, Hungary; University of Veterinary Medicine Budapest, Department of Food Hygiene, Budapest, 1078, Hungary

## Abstract

Antimicrobial resistance (AMR) is a global threat gaining more and more practical significance every year. The protection of bacteria against antimicrobials based on antimicrobial resistance genes (ARGs) developed in evolution. One of the essential clinical questions is the origin of ARGs of pathogen bacteria. Since the bacteria can share genetic components by horizontal gene transfer (HGT), all even non-pathogen bacteria may provide ARG to any pathogens when they became close physically. The bacteria of the human gut may make contact with bacteria entered into the body by food. The fermented food contains bacteria in high amount by its nature. Here we studied the diversity of ARG content by a unified metagenomic approach in various kefir and yoghurt products, in grain and isolated bacterial strains. We found numerous ARGs of commonly used fermenting bacteria with diversity characteristics in kefir and yoghurt samples. Even with the strictest filter restrictions we identified ARGs undermining the efficacy of aminocoumarin, aminoglycoside, carbapenem, cephalosporin, cephamycin, diaminopyrimidine, elfamycin, fluoroquinolone, fosfomycin, glycylcycline, lincosamides, macrolide, monobactam, nitrofuran, nitroimidazole, penam, penem, peptide, phenicol, rifamycin, tetracycline and triclosan. In the case of gene *lmrD*, we detected genetic environment providing mobility of this ARG. Our findings support that theory during the fermentation process the food ARG content can grow by the bacteria multiplication. Results presented suggest that starting culture strains of fermented food should be monitored and selected to decrease the ARG amount intake by nutrition.

## Introduction

Antimicrobial resistance (AMR) is a global threat gaining more and more practical significance every year. While antimicrobial resistance genes (ARGs) have been present ever since the appearance of the first living microorganisms as a part of their natural rivalry^1^, the real turning point came with the daily use of antibiotics in human and animal healthcare. Even though antibiotic compounds are not directly responsible for the genetic changes behind the appearance of antimicrobial resistance, they place selective pressure towards the amplification of individual bacterial ARGs. Identifying the sources of intake of bacteria carrying ARGs is nowadays a biomedical priority. Bacteria appear in the newborn body right by birth^2^, and later on, their invasion continues from the environment, from other humans and animals, or raw^3^ and processed^4^ food^5^. Bacteria reaching our gut through alimentation may share functional ARGs with horizontal gene transfer (HGT) either with saprophytes or with pathogens in their physical proximity. In the latter case, resistant or multi-resistant pathogenic strains may evolve, and therapeutic options for the treatment of bacterial infections may narrow. By the production and consumption of livestock products, the opportunity of establishing transfer connections between bacteria is provided as bacterial populations of different sets of resistance genes may meet. Thus, yoghurt and kefir, which are popular probiotic products commonly consumed without additional heat treatment, have a potential for providing a chance for such bacterial encounters. They are probiotic food with minor differences in their processing steps fermented with bacteria or with bacteria and fungi. They have both been present in the human diet for long and still stand their ground in today’s demanding, health-conscious society setting high nutritional standards. Nevertheless, besides the health benefits, the consumption of raw, probiotic food may have an adverse effect. Along with the multiplication of bacteria during the fermentation process, the bacterial resistome also grows. With the intake of raw products with probiotic strains, a higher possibility of HGT is provided in the human gut if the right triggers are present. Our study aimed to examine the diversity of ARG content by a unified metagenomic approach in various kefir and yoghurt products, in grain and isolated bacterial strains.

## Results

### Bacteriome

By taxon classification the number of reads aligning to bacterial genomes differed in the various samples (Fig 1/a). Two samples (k_g_04 and y_g_01 from bioproject PRJNA644779) contained ~ 20 million reads of bacterial origin. From bioproject PRJNA388572 sample k_p_15 had ~ 50 million bacterial reads, while k_p_14 contained more than 63 million. Excluding these four extremities the average bacterial read count of the metagenomic samples was 6.7 × 10^5^ (ranging between 7.3 × 10^4^ and 1.4 × 10^6^). The median sequencing depth of the strain k_s_01, k_s_02, k_s_03, k_s_04, k_s_05, k_s_06, k_s_07 was 46, 119, 115, 111, 6, 54, 108, respectively.

**Figure 1.**
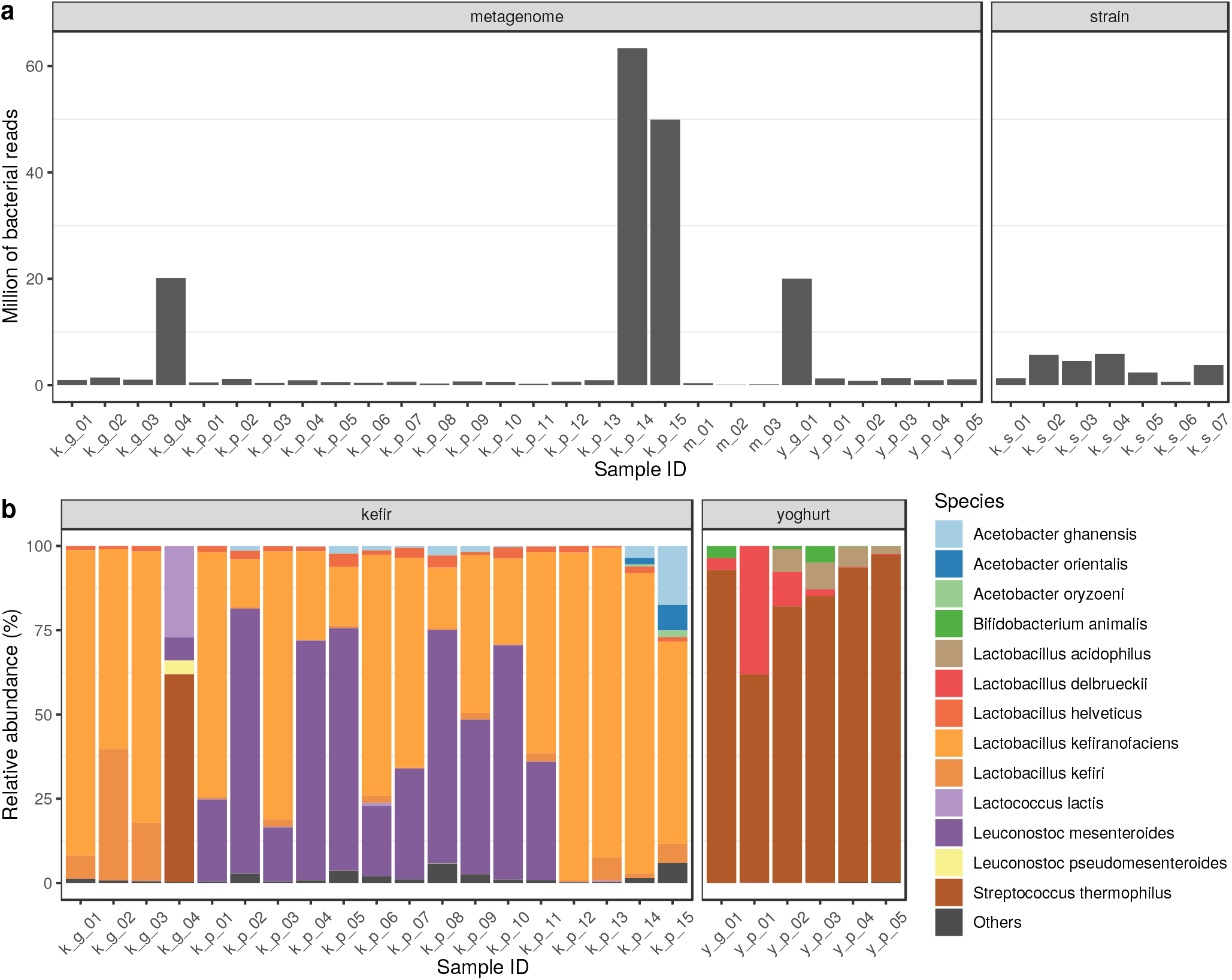
Bacterial content of the samples. **a** The number of reads classified bacterial by Kraken2 on the NCBI NT database. Metagenome includes the samples deriving from grains, milk or products. **b** Relative abundances of the most common bacterial species in grain and product samples.

**Figure 2.**
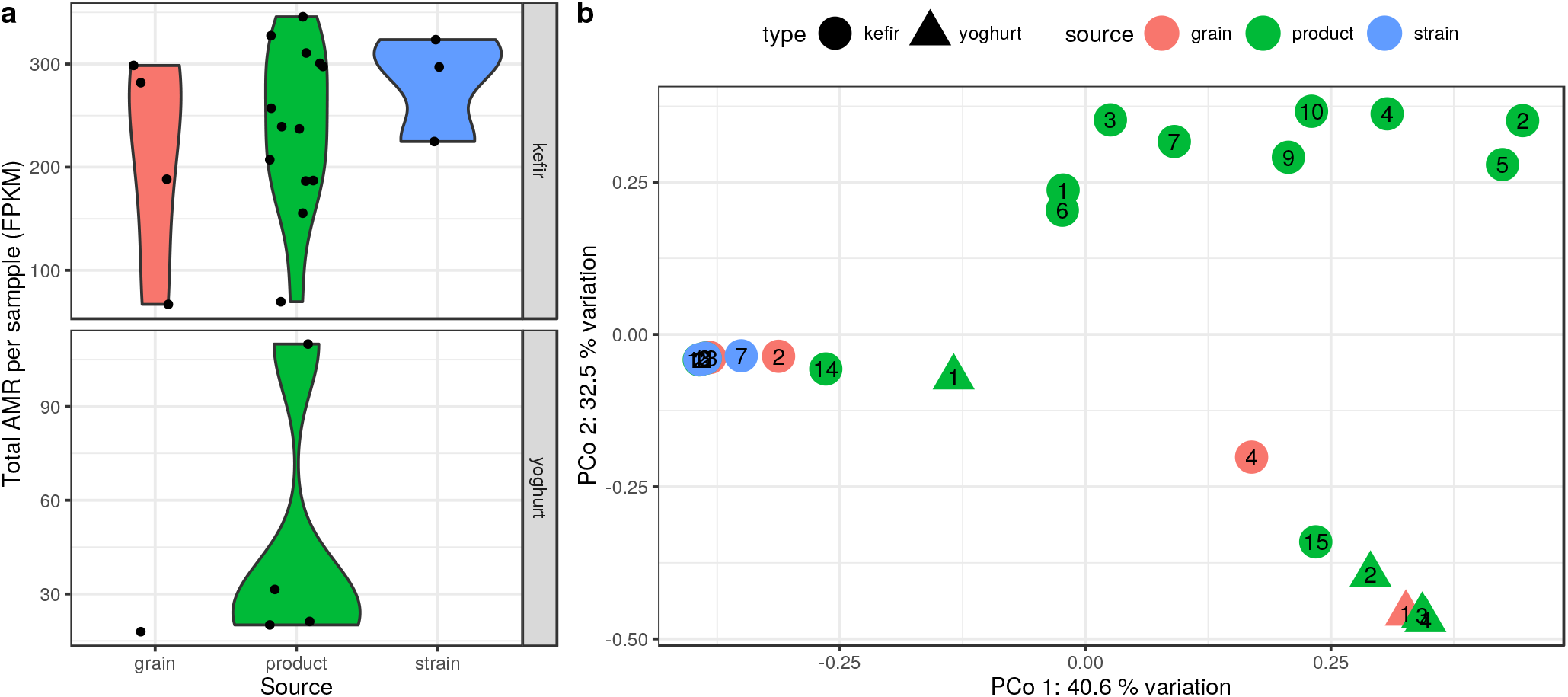
Antimicrobial resistance (AMR) abundance of the samples. **a** Violin plot represented distribution of the total AMR fragments per kilo base per million fragments (FPKM) per sample, grouped by type and source. The horizontally jittered dots represent the FPKM of the samples. **b** The AMR abundance diversity (*β*-diversity) of the samples. It is plotted on the first two axes of principal coordinate analysis (PCoA) performed on Bray-Curtis distance calculated using the relative abundances of contigs harboring ARGs. The symbols show the type, the colors the source, while the numbers correspond to the sequence number in the Sample ID.

Figure 1/b. demonstrates the relative abundance of dominant bacterial species identified in the samples. 99% of all bacteria identified was related to these species. In kefir grains dominant species were a *Lactobacillus kefiranofaciens* (57.7% ± 40.5%), *Lactobacillus kefiri (15.7%o* ± 17%), *Streptococcus thermophilus* (15.4% ± 30.8%), *Lactococcus lactis* (6.8% ± 13.5%), *Leuconostoc mesenteroides* (1.7% ± 3.4%), *Leuconostoc pseudomesenteroides* (1% ± 2%), *Lactobacillus helveticus* (1% ± 0.7%) in a descending order of their abundance rates. In products the most significant species overlapped the ones in the kefir grains with differences in their relative abundance rates (*L. kefiranofaciens* (55.4% ± 29%), *L. mesenteroides* (35.7% ± 30%), *Acetobacter ghanensis* (2.1% ±4.4%), *L. helveticus* (2.1% ± 1%), *L. kefiri* (1.8% ±2%), *Acetobacter orientalis* (0.6% ± 2%), *Acetobacter oryzoeni* (0.2% ± 0.5%)). The one yoghurt grain examined was dominated by *Streptococcus thermophilus* (92.8%), *Bifidobacterium animalis* (3.6%) and *Lactobacillus delbrueckii* (3.5%) while the core bacteriome of the product, yoghurt consisted of *S. thermophilus* (83.9% ± 13.8%), *L. delbrueckii* (10.1% ± 16.2%), *Lactobacillus acidophilus* (4.6% ± 3.3%), and *B. animalis* (1.2% ±2.1%).

### Resistome

According to our findings based on perfect and strict matches AMR gene abundances show a great diversity in various types and sources of samples. (Fig. 3/a). The highest AMG abundance was observed in the kefir strain samples (average: 282 fragments per kilo base per million fragments (FPKM), sd: 51.1) followed by the kefir product (240±78.6) and the kefir grains (209±106). The yoghurt samples had lower abundances, in the one grain FPKM was 17.9, while in the products we found 45.7±32.2.

**Figure 3.**
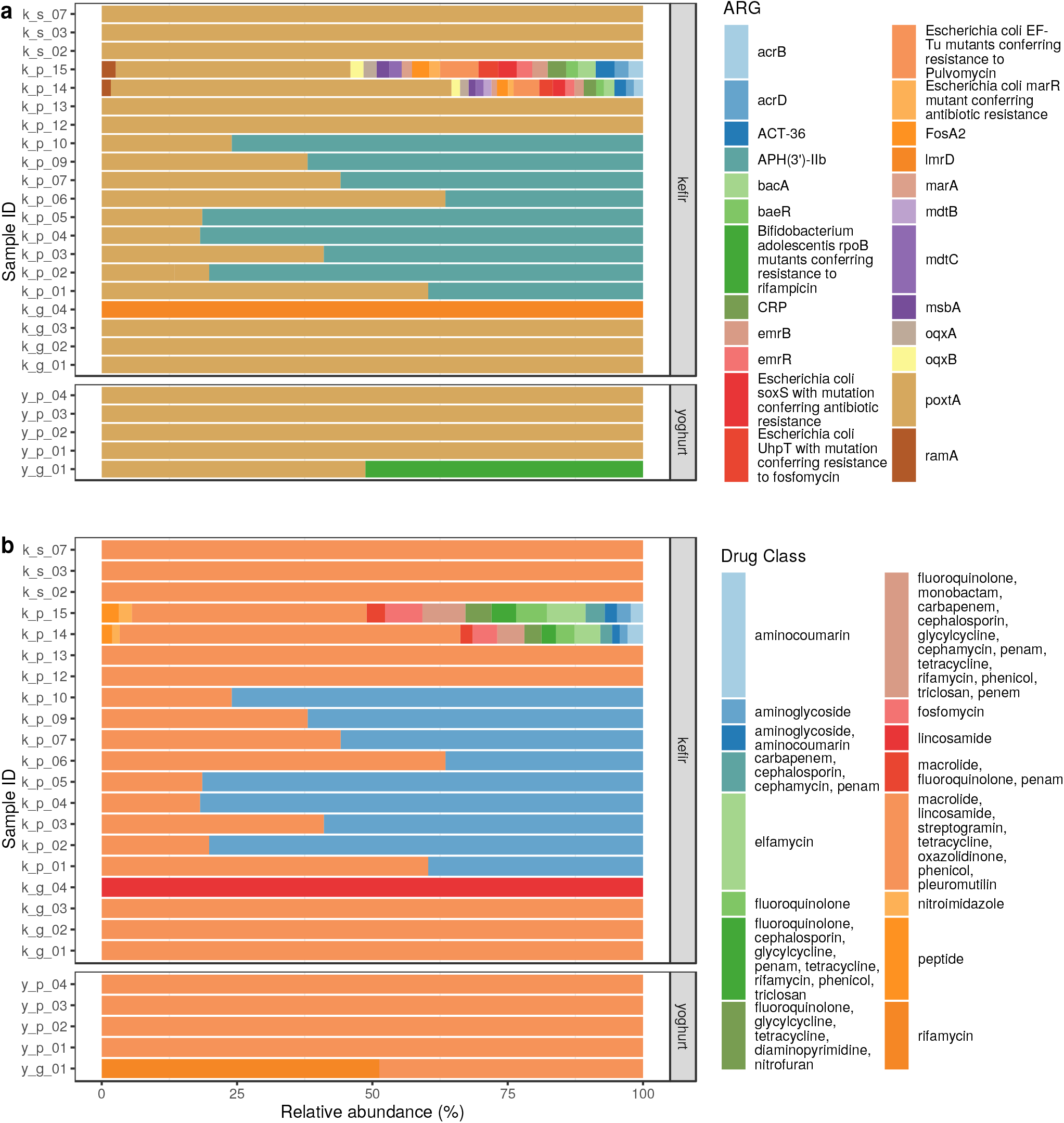
Antimicrobial resistance (AMR) abundance in kefir and yoghurt samples. **a** Relative abundance of AMR genes. ORFs having at least 60% length and 90% base sequence identity with the reference ARG sequence are shown. Kefir strains 1., 4., 5. and 6. are not shown as their ORFs did not meet these requirements. **b** Relative abundance of drug classes related to the ARGs identified in the samples.

**Figure 4.**
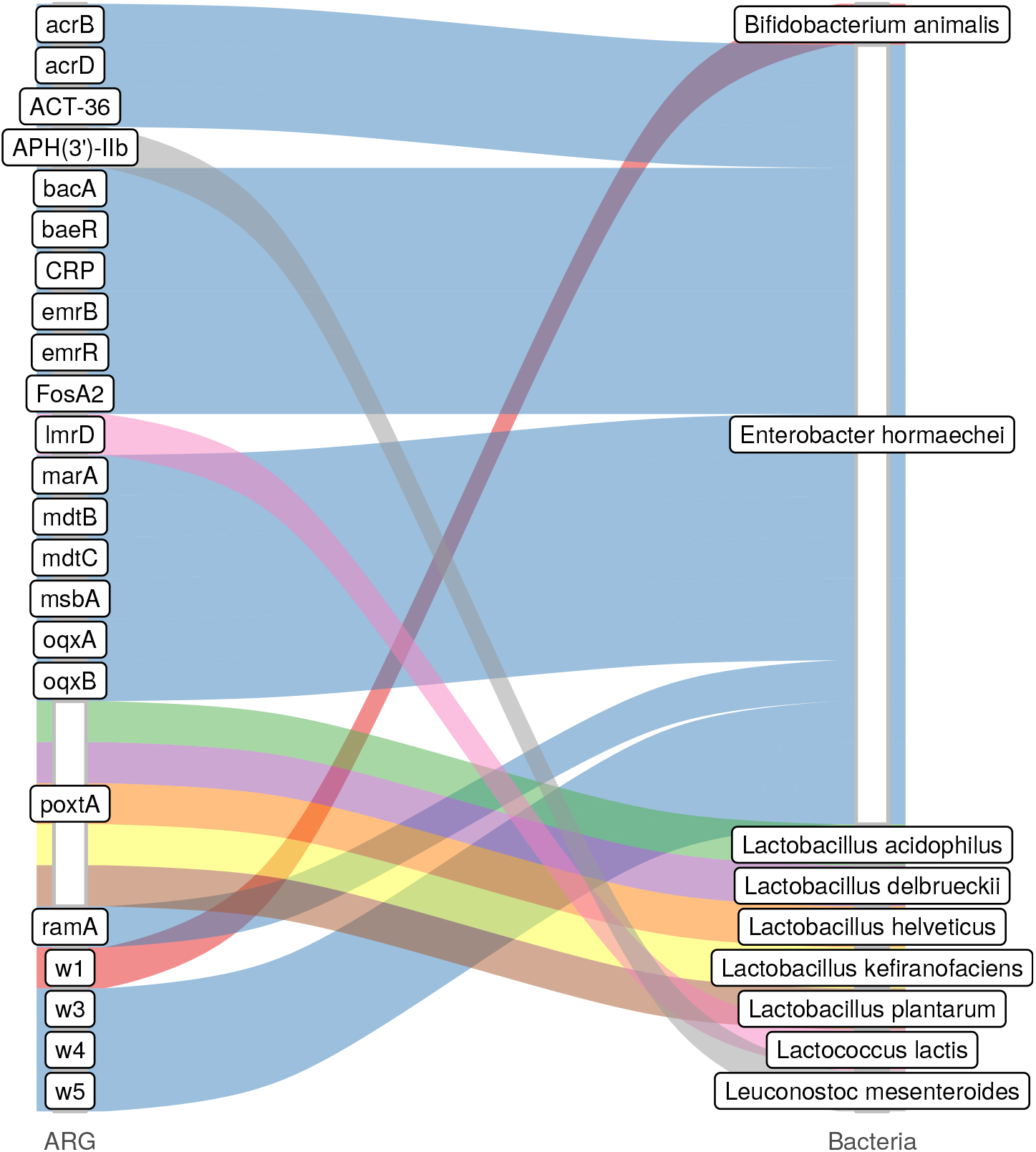
Identified ARGs and their most probable bacteria of origin The gene names that are too long have been abbreviated like w1: *Bifidobacterium adolescentis rpoB* mutants conferring resistance to rifampicin; w3: *Escherichia coli marR* mutant conferring antibiotic resistance; w4: *E. coli soxS* with mutation conferring antibiotic resistance; w5: *E. coli UhpT* with mutation conferring resistance to fosfomycin.

A Bray-Curtis distance based principal coordinate analysis (PCoA) was performed to gain insight into the dissimilarity of the sample ARG abundances (Fig. 2/b). With a permutational multivariate analysis of variance on the same distance matrix we found the type of the sample explains the 22.17% (p<0.001) of the dissimilarity among the sample resistomes. For the source grouping the same measure was 18.92% (p<0.001). Based on Fig. 2/b one might conclude that the strongest effect on the dissimilarity is the bioproject of origin, the analysis showed that it explains 35.56% (p<0.001) of the dissimilarity variances.

By kefir, we identified 22 ARGs in the product, 2 in the grain and 1 in the strain. By yoghurt there were 1 ARG in the product and 2 in the grain (Fig. 3/a). The relative abundances of antibiotic classes affected are shown in Fig. 3/b in each sample. By kefir ARGs identified in the product may help bacteria in the defense against aminocoumarins, aminoglycosides, carbapenems, cephalosporins, cephamycins, diaminopyrimidines, elfamycins, fluoroquinolones, fosfomycins, glycylcyclines, lincosamides, macrolides, monobactams, nitrofurans, nitroimidazoles, oxazolidinones, penams, penems, peptides, phenicols, pleuromutilins, rifamycins, streptogramins, tetracyclines and triclosan. Contigs containing these ARGs belonged to the genomes of *Enterobacter hormaechei* (genes: *acrB; acrD; ACT-36; bacA; baeR; CRP; emrB; emrR; Escherichia coli marR* mutant conferring antibiotic resistance; *E. coli soxS* with mutation conferring antibiotic resistance; *E. coli UhpT* with mutation conferring resistance to fosfomycin; *FosA2; marA; mdtB; mdtC; msbA; oqxA; oqxB; ramA), L. helveticus* (gene *poxtA), L. kefiranofaciens* (gene *poxtA*) and *L. mesenteroides* (gene *APH(3’)-IIb*). ARGs originating from the kefir grain may play a role in the appearance of resistance against lincosamides, macrolides, oxazolidinones, phenicols, pleuromutilins, streptogramins and tetracyclines and were found in the genomes of *L. kefiranofaciens* (gene *poxtA*) and *L. lactis* (gene *lmrD*). Gene *poxtA* deriving from kefir strains (*L. kefiranofaciens* and *L. plantarum)* confers resistance against lincosamides, macrolides, oxazolidinones, phenicols, pleuromutilins, streptogramins and tetracyclines. Genes found in yoghurt grains encoded resistance against lincosamides, macrolides, oxazolidinones, phenicols, pleuromutilins, rifamycins, streptogramins and tetracyclines, while the ARGs of the product itself may weaken the efficacy of lincosamides, macrolides, oxazolidinones, phenicols, pleuromutilins, streptogramins and tetracyclines. Contigs involving ARGs of the yoghurt product could have been connected to *L. acidophilus* (gene *poxtA*) and *L. delbrueckii* (gene *poxtA)*, while the ARGs of the grains aligned to the reference sequence of *B. animalis* (gene *Bifidobacterium adolescentis rpoB* mutants conferring resistance to rifampicin) and *L. delbrueckii* (gene *poxtA).*

Based on the ARG abundances the proportion of resistance mechanisms was calculated for each sample. In the kefir product samples the dominant mechanism of identified ARGs was the antibiotic target protection (50.73%), followed by antibiotic inactivation (45.45%), antibiotic efflux (2.03%), antibiotic target alteration (1.07%), antibiotic efflux; reduced permeability to antibiotic (0.32%), antibiotic target alteration; antibiotic efflux; reduced permeability to antibiotic (0.27%), antibiotic target alteration and antibiotic efflux (0.13%). In the kefir grain samples the main mechanisms were antibiotic target protection (91.98%) and antibiotic efflux (8.02%). In the kefir strains the antibiotic target protection was the only mechanism detected. Antibiotic target protection (100%) and antibiotic efflux (2.58%) may play a role in the appearance of resistance by yoghurt products. In the one yoghurt grain sample antibiotic target alteration; antibiotic target replacement (51.28%) and antibiotic target protection (48.72%) are the possible resistance mechanisms.

### Mobile elements

The results of mobile genetic element (MGE) domain coexisting analysis showed that the ARG *lmrD* in sample k_g_04 might be mobile since the contig cotaining the gene had a transposase ORF within the distance of 10 ORFs. By the PlasFlow^6^ tool there was not any identifiable ARG harboring contig that may be originated from plasmids.

### ARG abundance changes during kefir fermentation

According to the analysis of metagenomic data from the work of Walsh et al.^7^ ARG abundances change during the fermentation process (Fig. 5/a). In case of all three grains (Fr1, Ick, UK3) *APH(3’)-IIb* is present in the kefirs fermented for 24 h, while it is absent in all the other time phases except for sample UK3 after 8 h. The *poxtA* was detectable in all samples except the 8 h Fr1 kefir one. The abundance fold change of 24 h with respect to grain samples was 0.10, 0.59 and 0.26 for the starter culture Fr1, Ick and Uk3, respectively. Between the 8 h and 24 h samples for Ick kefir the *poxtA* abundance showed a 0.34-fold change, in the UK3 sequence this value was 0.62.

Contigs harboring the above listed ARGs were classified taxonomically (Fig. 5/c). All contigs having the gene *APH(3’)-IIb* were classified to *L. mesenteroides*. The *poxtA*-containing contigs were assigned to the reference genome of *L. helveticus* and *L. kefiranofaciens.*

**Figure 5.**
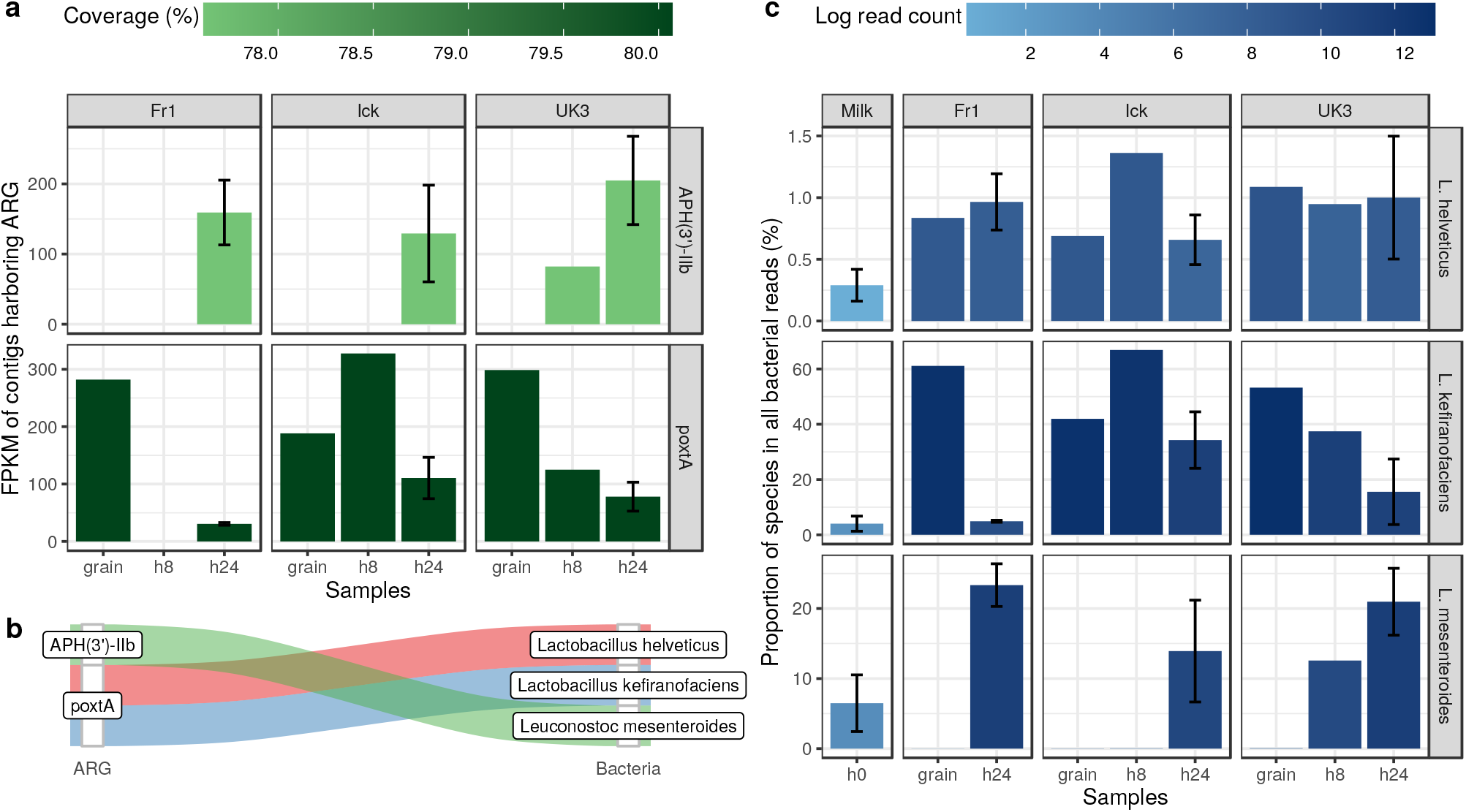
Changes during kefir fermentation. **a** Antimicrobial resistance gene (ARG) abundance expressed as fragments per kilo base per million fragments (FPKM) based on the aligment of bacterial reads to the ARG harboring contigs. **b** ARGs and their most likely origins. **c** Relative abundances of bacteria with a probable ARG content.

All bacterial reads were then aligned to the reference genomes of bacteria mentioned above and the hits were expressed as their proportion (Fig. 5/b). In contrast to *L. kefiranofaciens* that showed a decreasing tendency, an increase in time is observable by the relative abundances of *L. mesenteroides.* The proportion of reads assigned to *L. helveticus* shows no tendential increase or decrease in time. The increase of abundance of *APH(3’)-IIb* shows a positive association with the relative abundance of bacteria (*L. mesenteroides* detected to carry ARG harboring contigs. Similarly, *poxtA* abundance is decreasing with the relative abundance of *L. kefiranofaciens*.

### Excluding nudged hits

In order to set the alignment restrictions of ORFs to ARGs even stricter we may select for a subgroup of reference ARGs fitting the ORFs from the starting base position on. Thus, we do not permit nudging on the reference sequence by the alignment. With such a shrinkage we may reduce the number of detectable ARGs to four samples (Fig 6). Sample k_g_04 from bioproject PRJNA644779 contained an ARG against lincosamides while the gene found in sample y_g_01 is responsible for resistance against rifamycin. Contigs harboring these ARGs had the best alignment to *L. lactis* and *B. animalis*, respectively. Bioproject PRJNA388572 had two samples with similar matches, except for gene *mdtB* which appeared in full length in sample k_p_14 and was absent in k_p_15. This gene is responsible for aminocoumarin resistance. As some other ARGs of the sample also have the potential to confer resistance against this substance, the AMR profiles of the samples are the same, including aminocoumarin, aminoglycoside, carbapenem, cephalosporin, cephamycin, diaminopyrimidine, elfamycin, fluoroquinolone, fosfomycin, glycylcycline, macrolide, monobactam, nitrofuran, nitroimidazole, penam, penem, peptide, phenicol, rifamycin, tetracycline and triclosan. Comparing this list to the nudged results oxazolidinone, pleuromutilin and streptogramin resistance genes were absent. Contigs containing ARGs had the best alignment to the genome of *E. hormaechei* in both cases.^8^

**Figure 6.**
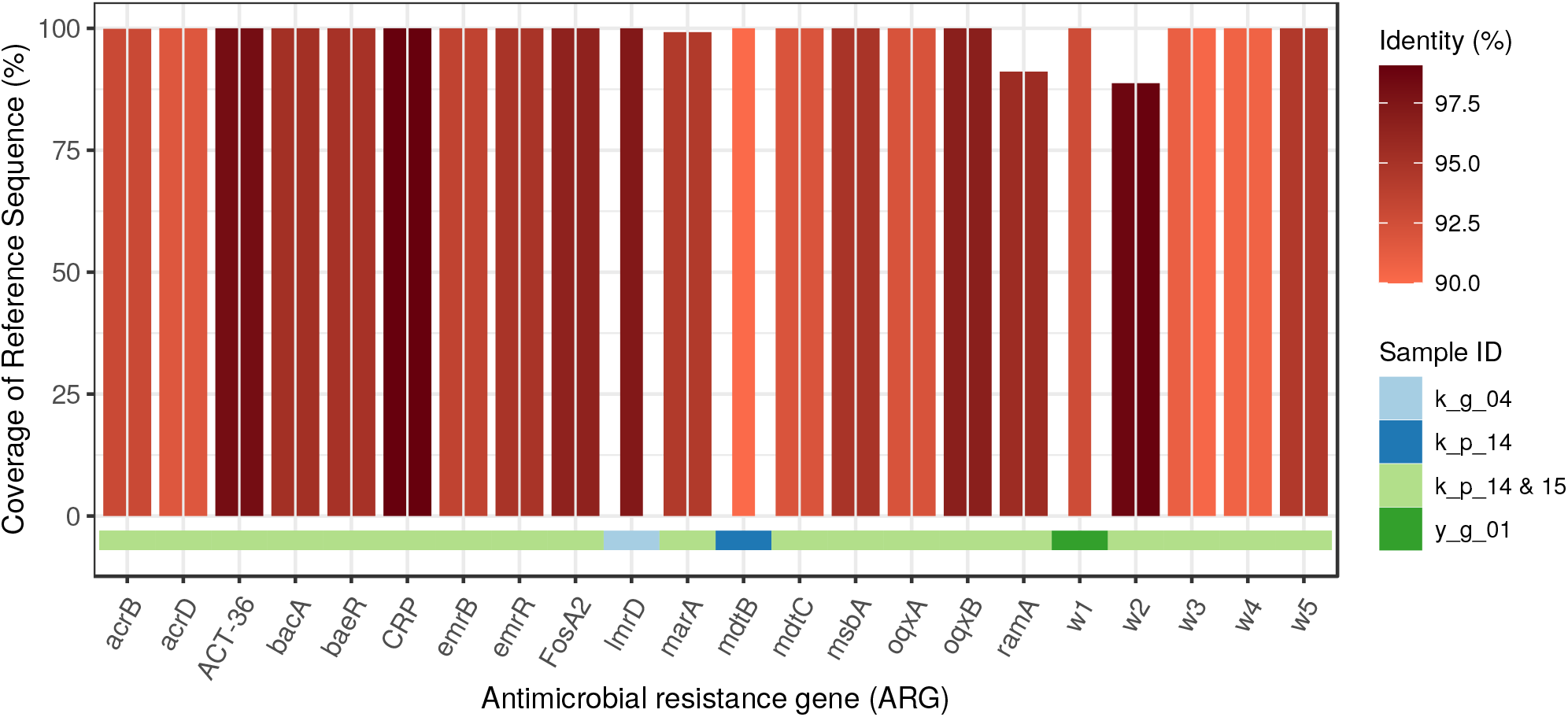
Identifed ARGs excluding nudged findings. The coverage and identity of detected open reading frames (ORFs) by antimicrobal resistance genes (ARGs). The ORF covered proportion of the reference ARG sequence and the identity % of predicted protein (color). The gene names that are too long have been abbreviated like w1: *Bifidobacterium adolescentis rpoB* mutants conferring resistance to rifampicin; w2: *Escherichia coli EF-Tu* mutants conferring resistance to Pulvomycin; w3: *E. coli marR* mutant conferring antibiotic resistance; w4: *E. coli soxS* with mutation conferring antibiotic resistance; w5: *E. coli UhpT* with mutation conferring resistance to fosfomycin.

All four samples of both bioprojects included at least 20 million bacterial reads in the assembly of the contigs. The other samples consisted of significantly less reads. Consequently, as Sims et al.^9^ found it is not possible to resolve whether an absence of a protein-coding gene, or a disruption of its open reading frame (ORF), represents a deficiency of the assembly or a real evolutionary gene loss. Examining the relationship among the number of bacterial reads and length of identified ARGs (including nudge) with a linear model we found that after each extra 100 000 read the coverage of reference gene raises 7% by the ORFs identified as ARGs (p<0.0001). By sample k_g_04, k_p_14, k_p_15 és a y_g_01 we randomly chose the average read number of the other samples (677 340) to reanalyze how much these results differ from the original ones executed on the full database. Contigs assembled contained one gene previously identified (excluding nudge), namely *lmrD* from sample k_g_04. ORFs previously identified as ARGs had a median coverage of 16.10% on the reference ARGs. In contrast, ORFs aligning to ARGs composed of the full read content of the four samples had a median coverage of 99.21%.

## Discussion

Studying ARGs that may enter the body by food, including fermented dairy products, can lead to critical health considerations. Both in bacterial and ARG diversity, there are observable characteristics in kefir and yoghurt. The ARG abundance is much higher in kefir than in yoghurt. One possible reason for this phenomenon could be the presence of fungi in kefir seed cultures. Since fungi may produce antibacterial toxins, bacteria having ARGs may gain a competitive advantage in the coexistence with fungi.

Each of the bacteria (*Bifidobacterium animalis*^10^, Enterobacter hormaechei^8,11^, Lactobacillus acidophilus^11,12^, Lactobacillus delbrueckii^11,12^, Lactobacillus helveticus^11,12^, Lactobacillus kefiranofaciens^11,12^, Lactobacillus plantarum^11,12,14^, *Lactococcus lactis*^11,12,15,16^, *Leuconostoc mesenteroides*^11,12,15,16^) obtained from the taxon classification of contigs containing ARGs is a widely used species in the production of fermented dairy products. Li et al.^17^ analyzed the ARG content of isolated bacteria from lactic acid bacterium drinks and yoghurts. They identified the gene *APH(3”)-III, APH(6’)-APH(2”), sul1, tet(M)* in *Lactobacillus bulgaricus* strains, while from *Streptococcus thermophilus* ones the gene *APH(3”)-II, sul1, sul2, strA, strB, tet(M)*. In our study, we found the *APH(3’)-IIb* gene belonging to the APH gene family, most probably originated from *L. mesenteroides.* Similarly, Carr et al.^18^ found a strong co-occurrence between *APH(3’)-Ia* and *L. mesenteroides* in China saliva samples. Further similarity between our and Carr et al.^18^ results is the *Lc. lactis* origin of *lmrD.* Guo et al.^19^ detected ARGs in *Lactobacillus* strains of fermented milk products. In strains of *Lactobacillus casei* gene *erm(B), gyrA, rpoB, vanE, vanX*, in *L. helveticus* strains gene *APH(3”)-III, dfrD, erm(B), gyrA, tet(W), vanX*, in *L. plantarum* isolates gene *erm(B)* and *vanX.* We found the *poxtA* gene associated with *L. helveticus* and *L. plantarum.* The *emrB* gene was identified by a contig from the genome of *E. hormaechei*.

During the milk fermentation process the bacteria of seed cultures (and the milk) multiply and dominate the beverage. If these bacteria harbours ARGs, then the amount of these genes will be increased in the final products. Based on data generated by Walsh et al.^7^ the bacteria *L. helveticus* and *L. kefiranofaciens* is the most probable origin of the contigs harbouring gene *poxtA*, while the gene *APH(3’)-IIb* containing sequences could have been stemmed from *L. mesenteroides*. In the original paper of Walsh et al.^7^ they found that under the fermentation the relative abundance of *L. kefiranofaciens* and *L. mesenteroides* decreased and increased by the time, respectively. Not surprisingly, in our reanalysis of the same data we found the same trends (Fig. 5/b). While Marsh et al.^20^ presented same changes of these species in kefir, Bao et al.^21^ showed opponent alterations in koumiss fermentation. Examining the changes in the ARG abundances (Fig. 5/a) and the relative abundances of their most probable bacteria of origin (Fig. 5/b) one may observe a positive association of them. With the increase of the relative abundance of *L. mesenteroides* the *APH(3’)-IIb* abundance rises, while lower *poxtA* abundance was observed by the decrease of the relative abundance of *L. kefiranofaciens*.

The two most abundant ARGs were *poxtA* and *APH(3’)-IIb*, the former present both in yoghurt and kefir samples. *PoxtA* (phenicol-oxazolidine-tetracycline resistance), an abundant ARG in Gram-positives confers resistance to a wide range of highly important antibiotics. ABC-F class ATP binding ribosomal protection protein encoded by this gene is present mainly in Enterococci and Staphylococci. It was identified in an methicillin-resistant *Staphylococcus aureus* (MRSA) strain that showed increased MIC to linezolid, a member of the oxazolidine class of ABs.^22^ The study highlighted that Staphylococci, Enterococci and interestingly, *Pediococcus acidilactici* harboring the gene are all of animal origin, and can be spread horizontally via mobile genetic elements. In line with other papers^23^ the study suggests that phenicols and other antiribosomal agents used in veterinary medicine might have played a role in the selection of *poxtA.* This was also confirmed by Elghaieb et al.^24^ who identified the gene in cow milk and animal wastewater. As oxazolidines are prohibited in food animals and phenicols are not permitted in dairy cattle in Europe, the source of these genes in Hungarian samples remains to be elucidated. *Pseudomonas aeruginosa* harbors an array of aminoglycoside-modifying genes, altering the drug by acetylation, adenylation or phosphorylation (APH). The presence of *APH(3’)-IIb* in kefir samples is deliberately worrying as aminoglycoside 3’-phosphotransferases can mediate high level resistance against several aminoglycosides. These genes might be plasmid-borne or chromosomally encoded; *APH(3’)-IIb* is the latter, but a transposon-mediated mechanism has been suggested to be responsible for spreading the resistance genes.^25,26^ As the gene was almost exclusively described in *P. aeruginosa* previously, and the likely origin was *L. mesenteroides* in our study, the routes of resistance gene transfer related to this gene needs to be further investigated. Although penicillins and cephalosporins are the most frequently used antibiotics in dairy cows, interestingly, the abundance of ARGs facilitating resistance against β-lactams are rather lacking. This phenomenon, together with the ARGs related to unused antibiotics in veterinary dairy medicine, raises the suspicion that the source of the abundant ARGs might not be a direct consequence of antibiotic use in dairy cattle.

Bacteria enter the digestive tract with food, what opens an opportunity to get close to other non-pathogenic and pathogenic bacteria. One of the main prerequisite of horizontal gene transfer (HGT) processes is the physical closeness among the participating bacteria. Through gene transfer various genes, including ARGs can be exchanged by bacteria. If a certain ARG is placed on a mobile DNA-sequence, then the probability of HGT is higher. We found only one gene, the lmrD in sample k_g_04 that might be mobile due to its genomic environment.

Antibiotic resistance and multi drug-resistant bacteria are a significant global public health threat.^27^ Infections caused by drug-resistant bacteria are causing major morbidity and mortality and increase the cost of health care when compared to infections by non-resistant strains of the same bacteria. WHO identified families of multi-drug-resistant bacteria, the so called “priority pathogens” classifying them into critical, high and medium risks based on their threat level to the human health. These bacteria are resistant to several antibiotics genes identified in our study highlighting the practical importance of our findings. Even with the strictest filter restrictions we identified ARGs undermining the efficacy of aminocoumarin, aminoglycoside, carbapenem, cephalosporin, cephamycin, diaminopyrimidine, elfamycin, fluoroquinolone, fosfomycin, glycylcycline, lincosamides, macrolide, monobactam, nitrofuran, nitroimidazole, penam, penem, peptide, phenicol, rifamycin, tetracycline and triclosan. These findings raises clinical considerations. Priority 1 (“critical”) class pathogens, *Acinetobacter baumannii, Pseudomonas aeruginosa*, and the *Enterobacteriaceae* are resistant to carbapenems, the best available antibiotics reserved as last resort treatments for treating multi-drug-resistant bacteria.^27^ Carbapenems possess the broadest spectrum of antimicrobial activity, and often reserved for the most severe infections and used as ultimum refugium agents. The common indications for treatment with a carbapenem (imipenem or meropenem) are urinary infections resistant to other antibiotics, treatment of necrotizing pancreatitis^28^, treatment of severe nosocomial infections including hospital-acquired pneumonia, intraabdominal infections, bacterial soft-tissue infections, febrile neutropenia. Tigecycline, a recently developed third-generation tetracycline antibiotic belonging to the glycylcycline class is one of the few therapeutic options for treating infections caused by *A. baumannii* and most carbapenem-resistant bacteria. Indications include hospital-acquired pneumonia and blood-stream infections (BSI) caused by *Klebsiella pneumoniae*^29,30^ and nosocomial urinary tract, pulmonary, abdominal, surgical site, and bloodstream infections caused by carbapenem-resistant *Enterobacteriaceae* (CRE).^31^ Another class of AMR identified in our study were fluoroquinolones. These antibiotics, for example ciprofloxacin, ofloxacin and levofloxacin are used for treating urinary tract and respiratory tract infections, bone and joint infections and gonorrhea. Emerging fluoroquinolone resistance in *Campylobacter* strains which are the leading cause of bacterial gastroenteritis in the world, is a significant public health concern similarly to the rising incidence of fluoroquinolone resistant cases of typhoid fever and invasive non-typhoidal *Salmonella* (iNTS) infections. We have also identified genes coding cephalosporin resistance in our samples. Cephalosoprins belong to the most frequently used antibiotics. Whereas first and second generation cephalosporins are effective against Gram-positive bacteria; intravenous third generation cephalosporins (eg. ceftriaxone) are more potent against Gram-negative bacteria and frequently used in cholecystitis, spontaneous bacterial peritonitis or as a preventive measure in acute gastrointestinal hemorrhage.^32^ Resistance to extended-spectrum cephalosporins (ESCs), for example cefixime and ceftriaxone and to fluoroquinolones is causing a rising incidence of treatment failures in gonorrhea triggering several action plans due to the large burden of gonorrhea globally. Due to recent emergence of multi-resistant Gram-negative bacteria (extended-spectrum-lactamase (ESBL)-producing *Enterobacteriaceae*, *Pseudomonas* and *Acinetobacter* spp) aminoglycosides previously neglected due to issues about their tocixicity have recently been re-evaluated in nosocomial urinary tract and respiratory infections.^33^ In our study we have found genes of lincosamide resistence. One of the most widely utilized lincosamide is the semisynthetic clindamycin used for BSI, intra-abdominal, gynecological infections and bone and joint infections.^34^ Clindamycin is also used in the dermatology in the treatment acne vulgaris and hidradenitis suppurativa and in prevention of surgical-site infections (SSIs) in patients allergic to penicillin.^35^ Furthermore, it is utilized for endocarditis prophylaxis in dentistry.^36^ Macrolide antibiotics have excellent absorption from the gastrointestinal tract and few side effects. Clarythromycin is still considered as a member of first-line treatment protocol for *Helicobacter pylori* eradication in areas with low resistance to clarythromycin^37^. Azithromycin can contribute to resolution of acute infections by immunomodulatory effects^38^. It is frequently used for the treatment of acute watery or febrile diarrhea and dysentery syndrome^39^. Once commonly used, nowadays rarely administered tetracyclin has been recently rediscovered, as a component of *H. pylori* eradication regimen, partly due to increasing rate of resistance to other antibiotics (including the above-mentioned clarythromycin)^37^. Resistance to macrolides could hamper treatment of toxoplasmosis in pregnancy^40^ and urogenital infections.^41^ as well. Cloramphenicole is listed on WHO Model List of Essential Medicines as an “access group antibiotic” which are the first and second choices for empirical treatment of 21 common clinical syndromes.^27^ Cloramphenicole is used in bacterial conjunctivitis, meningococcal sepsis and Rickettsial disease.^42,43^ Standard treatment of tuberculosis consists a a combination of isoniazid, rifampicin, ethambutol, and pyrazinamide. Increasing prevalence of rifamycin-resistant tuberculosis is a growing problem globally.^44^

As ARGs reaching the human body may originate from fermented dairy products, further examinations would be worthwhile to clarify the details and understand the practical medical significance. For this, it would be expedient to analyze the samples of starter cultures and final products and register the results at time points previously set throughout the fermentation period. According to our findings, sequencing depth plays a significant role in the coverage of ORFs identified as ARGs, thus involving at least 20 million clusters is recommended by similar studies. The samples we examined and the studies found in the literature^3–5,17,19^ confirm the hypothesis that foods of animal origin may contain significant amounts of diverse AMGs. The reasons for the appearance of the AMGs are complex and difficult-to-control. As sequencing techniques become cheaper, in addition to further strict control of antibiotics used in animal husbandry, regular genetic monitoring of products of animal origin, including in our case starter cultures, should be considered.

## Methods

### Data

The details of analyzed samples are listed in Table 1. One kefir and one yoghurt starter culture was shotgun sequenced (PRJNA644779) by the authors. The further short read datasets were obtained from NCBI SRA repository. A query was performed in SRA to find kefir or yoghurt related shotgun sequenced samples. As a result of searching more 33 datasets originated from 8 BioProjects were selected to the study. Except the samples of BioProjects PRJEB15432 all others came from paired end runs. The downloaded short reads were originated from BioSamples of kefir grain (n=4), kefir product (n=15), kefir strain (n=7), yoghurt grain (n=1) and yoghurt product (n=5). In the unified names of the samples the first character corresponds to the type of the sample (k and y, kefir and yoghurt, respectively), the second character comes from the first letter of the source (g, p and s for grain, product and strain, respectively), while the last tag is a sequence number. From the collected projects for the PRJNA222257^45^, PRJEB15432^7^ and PRJEB30083^46^ peer-reviewed publication is available. For the other samples the only accessible metadata was the attributes in SRA. In PRJEB15432 Walsh et al.^7^ followed the microbial changes during the fermentation process of kefir. They used full-fat pasteurized milk inoculated by three different grains (Fr1, Ick, and UK3 from France, Ireland and United Kingdom, respectively). The pasteurized milk (with three replications) and grains (without replication) were sampled at hour 0. In the fermentation from kefir at hour 8 (without replication) and hour 24 (with three replications) further specimens were taken.

**Table 1.** The list of analyzed samples obtained from NCBI SRA. In the unified names of the samples the first character corresponds to the type of the sample (k and y, kefir and yoghurt, respectively), the second character comes from the first letter of the source (g, p and s for grain, product and strain, respectively), while the last tag is a sequence number. The last column shows the available attribute data about the biosamples.

| Sample ID | BioProject | Run | Type | Source | Sample |
| --- | --- | --- | --- | --- | --- |
| k_g_01 | PRJEB15432 | ERR1653138 | kefir | grain | Fr1 grain |
| k_g_02 | PRJEB15432 | ERR1653139 | kefir | grain | Ick grain |
| k_g_03 | PRJEB15432 | ERR1653140 | kefir | grain | UK3 grain |
| k_g_04 | PRJNA644779 | SRR12171332 | kefir | grain | kefir seed culture |
| k_p_01 | PRJEB15432 | ERR1653129 | kefir | product | UK3, 8 hours |
| k_p_02 | PRJEB15432 | ERR1653130 | kefir | product | Fr1, 24 hours (replicate 2) |
| k_p_03 | PRJEB15432 | ERR1653131 | kefir | product | Ick, 24 hours (replicate 2) |
| k_p_04 | PRJEB15432 | ERR1653132 | kefir | product | UK3, 24 hours (replicate 2) |
| k_p_05 | PRJEB15432 | ERR1653135 | kefir | product | Fr1, 24 hours (replicate 3) |
| k_p_06 | PRJEB15432 | ERR1653136 | kefir | product | Ick, 24 hours (replicate 3) |
| k_p_07 | PRJEB15432 | ERR1653137 | kefir | product | UK3, 24 hours (replicate 3) |
| k_p_08 | PRJEB15432 | ERR1653141 | kefir | product | Fr1, 24 hours (replicate 1) |
| k_p_09 | PRJEB15432 | ERR1653142 | kefir | product | Ick, 24 hours (replicate 1) |
| k_p_10 | PRJEB15432 | ERR1653143 | kefir | product | UK3, 24 hours (replicate 1) |
| k_p_11 | PRJEB15432 | ERR1653145 | kefir | product | Fr1, 8 hours |
| k_p_12 | PRJEB15432 | ERR1653146 | kefir | product | Ick, 8 hours |
| k_p_13 | PRJNA288044 | SRR2082409 | kefir | product | KEFIR.shotgun |
| k_p_14 | PRJNA388572 | SRR7287342 | kefir | product | Metagenome from probiotic beverage K03 |
| k_p_15 | PRJNA388572 | SRR8282406 | kefir | product | Metagenome from probiotic beverage K02 |
| k_s_01 | PRJDB4955 | DRR064132 | kefir | strain | *Lactobacillus parakefiri* JCM 8573 |
| k_s_02 | PRJNA222257 | SRR1151211 | kefir | strain | *Lactobacillus kefirano-faciens subsp. kefirano-faciens* DSM 5016 |
| k_s_03 | PRJNA222257 | SRR1151212 | kefir | strain | *Lactobacillus kefirano-faciens subsp. kefir-granum* DSM 10550 |
| k_s_04 | PRJNA222257 | SRR1151213 | kefir | strain | *Lactobacillus kefiri* DSM 20587 |
| k_s_05 | PRJNA222257 | SRR1151226 | kefir | strain | *Lactobacillus parakefiri* DSM 10551 |
| k_s_06 | PRJNA635855 | SRR11965732 | kefir | strain | *Acetobacter syzygii* str. K03D05 |
| k_s_07 | PRJNA635872 | SRR11966381 | kefir | strain | *Lactobacillus plantarum* K03D08 |
| m_01 | PRJEB15432 | ERR1653133 | milk | milk | 0 hours (replicate 1) |
| m_02 | PRJEB15432 | ERR1653134 | milk | milk | 0 hours (replicate 2) |
| m_03 | PRJEB15432 | ERR1653144 | milk | milk | 0 hours (replicate 3) |
| y_g_01 | PRJNA644779 | SRR12171305 | yoghurt | grain | yoghurt seed culture |
| y_p_01 | PRJEB30083 | ERR2982980 | yoghurt | product | Yoghurt-A |
| y_p_02 | PRJEB30083 | ERR2982981 | yoghurt | product | Yoghurt-B |
| y_p_03 | PRJEB30083 | ERR2982982 | yoghurt | product | Yoghurt-C |
| y_p_04 | PRJEB30083 | ERR2982983 | yoghurt | product | Yoghurt-D |
| y_p_05 | PRJEB30083 | ERR2982984 | yoghurt | product | Yoghurt-E |

### DNA extraction and metagenomics library preparation for PRJNA644779

Total metagenome DNA of kefir (k_g_04) and yoghurt (y_g_01) samples were extracted by using the UltraClean Microbial DNA Isolation kit from MoBio Laboratories. The quality of the isolated total metagenome DNA was checked using an Agilent Tapestation 2200 instrument. The DNA samples were used for in vitro fragment library preparation. In vitro fragment libraries were prepared using the NEBNext Ultra II DNA Library Prep Kit for Illumina. Paired-end fragment reads were generated on an Illumina NextSeq sequencer using TG NextSeq 500/550 High Output Kit v2 (300 cycles). Read numbers were the following: 22 044 496 and 20 895 112 for kefir and yoghurt, respectively. Primary data analysis (base-calling) was carried out with Bbcl2fastq software (v2.17.1.14, Illumina).

### Bioinformatic and statistical analysis

Quality based filtering and trimming was performed by Trimmomatic^47^, using 15 as a quality threshold. Only reads longer than 50 bp were retained. The remaining reads were taxonomically classified using Kraken2 (*k* = 35)^48^ with the NCBI non-redundant nucleotide database^49^ with two different confidence setting. The first run was performed with the default settings to select all possible bacterial reads. In following taxon classification the `-confidence 0.5` parameter was set up to get more precise species level hits. The taxon classification data was managed in R^50^ using functions of package phyloseq^51^ and microbiome^52^. For further analysis, the reads assigned to Bacteria was used only^53^. By MEGAHIT (v1.2.9)^54^ with default settings the preprocessed bacterial origin reads were assembled to contigs. From the contigs all possible open reading frames (ORFs) were gathered by Prodigal^55^. The protein translated ORFs were aligned to the ARGs of database CARD v.3.0.9^56,57^ by Resistance Gene Identifier (RGI, v5.1.0) with Diamond^58^. The ORFs classified as perfect or strict were further filtered with 90% identity and 60% coverage. The findings were presented including and excluding the nudged hits as well. Contigs harbouring ARG identified by RGI with classified by Kraken2 with the NCBI RefSeq^59^ complete bacterial genomes database. Following Hendriksen at al.^1^ the ARG abundance was expressed with fragments per kilo base per million fragments (FPKM)^60^ of of contigs containing ARGs. For the ith contig *FPKM_i_* = *q_j_*(*l_i_* × *Q*) × 10^6^, where *q_i_* is the number of reads that mapped to the contig, *l_i_* is the length of contig and *Q* is the total number of mapped reads. To calculate *q* values all bacterial reads were aligned to the contigs by Bowtie2^61^ with parameter of `-very-sensitive-local`. To identify possible further mobile genetic element (MGE) homologs the predicted protein sequences of contigs were scanned by HMMER^62^ against data of PFAM v32^63^ and TnpPred^64^. Following Saenz et al.^53^ from the hits with lower than E 10^−5^ the best was assigned to each predicted protein within the distance of 10 ORFs. The MGE domains coexisting with ARGs were categorized as phage integrase, resolvase, transposase or transposon. The plasmid origin probability of the contigs was estimated by PlasFlow v.1.1^6^. According to ARG abundance of the samples a dissimilarity matrix was calculated using the Bray-Curtis index^65^ with package vegan^66^. With the same library using this matrix a permutational multivariate analysis of variance was applied to quantify the associations between the dissimilarity and independent variables (type, source, BioProject). For the visualization of the sample distances based on this matrix a principal coordinate analysis (PCoA) was performed with package ape^67^. The relationship between the detected ORF length and the sequencing depth was explored by linear model. All analysis and plotting was done in R-environment^50^.

## Acknowledgements

The project is supported by the European Union and co-financed by the European Social Fund (No. EFOP-3.6.3-VEKOP-16-2017-00005) and has received funding from the European Union’s Horizon 2020 research and innovation program under Grant Agreement No. 874735 (VEO).

## Author contributions statement

NS takes responsibility for the integrity of the data and the accuracy of the data analysis. AGT, IC, LM and NS conceived of the study concept. GM and NS performed the sample collection and procedures. AGT, NS and SAN participated in the bioinformatic and statistical analysis. AD, AGT, AJ, AVP, GM, GS, KB and NS participated in the drafting of the manuscript. AD, AGT, AJ, AVP, GM, GS, IC, KB and NS carried out the critical revision of the manuscript for important intellectual content. All authors read and approved the final manuscript.

## Additional information

### Availability of data and material

The short read data of sample data are publicly available and can be accessed through the PRJDB4955, PRJEB15432, PRJEB30083, PRJNA222257, PRJNA288044, PRJNA388572, PRJNA635855, PRJNA635872, PRJNA644779 from the NCBI Sequence Read Archive (SRA).

### Ethics approval and consent to participate

Not applicable.

### Consent for publication

Not applicable.

